# Modulation of metabolic, inflammatory, fibrotic and cell death pathways by Resmetirom in metabolic dysfunction-associated steatohepatitis (MASH): A transcriptomic profiling study

**DOI:** 10.1101/2025.11.26.690715

**Authors:** Chen-yang He, Zhi-hua Wang, Jian-ping Weng, Suo-wen Xu

## Abstract

Metabolic dysfunction–associated steatohepatitis (MASH) is a growing global burden with limited therapeutic options, and the mechanisms driving disease regression remain incompletely understood. Resmetirom, a selective thyroid hormone receptor-β (THR-β) agonist, has been approved for MASH, yet its molecular actions require further clarification. In this study, we employed a diet-induced mouse model of MASH using fructose–palmitate–cholesterol (FPC) feeding and evaluated the therapeutic efficacy of Resmetirom. Resmetirom (5 mg/kg/day) treatment significantly improved systemic and hepatic lipid profiles. Histological analyses demonstrated marked attenuation of hepatic steatosis, inflammation, and fibrosis, accompanied by reduced hepatic cell death. Transcriptomic profiling of liver tissues further revealed a robust Resmetirom-induced reprogramming of metabolic and inflammatory pathways, with prominent downregulation of gene networks governing lipid accumulation, immune activation, extracellular matrix remodeling, and multiple forms of programmed cell death. Collectively, these findings provide mechanistic insight into the multifaceted actions of Resmetirom and establish a comprehensive preclinical framework for understanding how THR-β activation ameliorates key pathological hallmarks of MASH. This work supports the therapeutic potential of Resmetirom and highlights critical molecular pathways that may inform future combination strategies for advanced metabolic liver disease.

## Introduction

Metabolic dysfunction–associated steatohepatitis (MASH) has emerged as a major contributor to chronic liver disease worldwide. Being categorized under metabolic dysfunction-associated steatotic liver disease (MASLD), MASH arises from simple steatosis in the context of metabolic dysfunction and can progress to advanced fibrosis, cirrhosis, and hepatocellular carcinoma^[1, 2]^. Closely linked to obesity, insulin resistance, and other components of metabolic syndrome, MASH has become one of the leading causes of liver-related morbidity and mortality globally^[3-6]^.

Resmetirom, a selective thyroid hormone receptor-β (THR-β) agonist, is the first drug approved by the U.S. Food and Drug Administration (FDA) for the treatment of MASH^[7, 8]^. Clinical trials have demonstrated its efficacy in reducing hepatic inflammation and fibrosis^[9, 10]^. However, the precise mechanisms of action underlying its hepatoprotective actions remain largely unclear. In particular, whether altered thyroid hormone receptor signaling contributes to disease regression and how Resmetirom influences key pathological processes such as hepatic steatosis, inflammation, fibrosis, and hepatic cell death are not fully defined. In this study, we employed a fructose–palmitate–cholesterol (FPC) dietary mouse model that recapitulates key features of human MASH, including steatosis, metabolic inflammation, collagen deposition, and hepatocyte death. By integrating histological, biochemical, and transcriptomic analyses, we sought to define the molecular mechanisms through which Resmetirom exerts its hepatoprotective effects. In particular, we focused on delineating genome-wide transcriptional changes induced by THR-β activation and identifying the specific metabolic, inflammatory, and cell death pathways that may mediate disease improvement. This integrated approach provides new mechanistic insight into how Resmetirom modulates the complex pathogenesis of MASH.

## Materials and methods

### Animal Models and Treatments

Six-week-old male C57BL/6J mice (#N000013, GemPharmatech) were purchased and acclimated for one week before the start of experiments. Mice were housed in standard cages under specific pathogen-free (SPF) conditions, with a 12-hour light/12-hour dark cycle at 22 ± 2°C, and had free access to food and water. To induce metabolic dysfunction-associated steatohepatitis (MASH), mice were fed a high-fructose, high-palmitate, and high-cholesterol diet (FPC diet; #TD.160785, Harlen Teklad) daily with a solution containing glucose and fructose (prepared by dissolving 23.1 g of glucose and 18.9 g of fructose in 1 L of sterile water) for a total of 12 weeks^[11]^. From week 4 to week 12, mice were randomly assigned to receive daily intraperitoneal injections of either Resmetirom (5 mg/kg/day) or a vehicle solution containing 0.5% carboxymethyl cellulose-sodium (#C9481; Sigma-Aldrich) and 0.4% Tween-80 (#A600562; Sangon Biotech). Mice were monitored closely, and body weight was recorded weekly throughout the feeding and treatment period. Experimental groups were designed to allow comparison between FPC-fed mice treated with Resmetirom and vehicle controls. All procedures were performed in accordance with institutional guidelines and approved by the Animal Ethics Committee of The First Affiliated Hospital of USTC (Approval No. 2025-N(A)-180).

### Histological Analysis of Liver Tissues

At the end of the experiment, mice were euthanized and livers were perfused with phosphate-buffered saline (PBS) to remove remaining blood. Entire livers were then excised, and anatomically matched sections were collected for further analyses. Tissue samples were fixed in 4% paraformaldehyde at 4°C overnight, followed by cryoprotection in 30% sucrose solution at 4°C until the tissues sank. For lipid detection, Fixed tissues were embedded in OCT compound (#4583, SAKURA) and cryosectioned at 10 μm thickness. Histological staining was performed following standard protocols. Hepatic lipid accumulation was assessed using Nile Red (#N3013, Sigma-Aldrich) and BODIPY (#793728, Sigma-Aldrich) staining^[12]^. Sections were stained with hematoxylin and eosin (H&E; #G1076, Servicebio) to assess general tissue morphology and with Sirius red (#G1427, Servicebio) to evaluate fibrosis. All staining procedures were performed following established protocols and the manufacturers’ instructions, ensuring consistent and reproducible results across samples.

### Serum Collection and Biochemical Analysis

At the end of the 12-week feeding period, mice were euthanized via intraperitoneal injection of sodium pentobarbital. Blood samples were collected and allowed to clot at room temperature for 3–4 hours. Following coagulation, samples were centrifuged at 8,000 rpm for 10 minutes at 4°C to obtain serum. Serum was subsequently analyzed for lipid profiles and liver function markers using commercial assay kits (Rayto, Shenzhen, China). Parameters measured included triglycerides (TG; #S03027), total cholesterol (TC; #S03042), high-density lipoprotein cholesterol (HDL-C; #S03025), low-density lipoprotein cholesterol (LDL-C; #S03029), aspartate aminotransferase (AST; #S03040), and alanine aminotransferase (ALT; #S03030), following the manufacturers’ protocols.

### Hepatic Lipid Quantification

Approximately 25 mg of liver tissue was carefully excised and weighed for lipid analysis. Hepatic triglyceride and total cholesterol levels were measured using commercial assay kits (LabAssay Triglyceride, #290-63701; LabAssay Cholesterol, #294-65801, FUJIFILM Wako) according to the manufacturers’ instructions.

### Immunofluorescence Staining

Frozen liver sections were rinsed with phosphate-buffered saline (PBS) for 10 minutes to remove residual OCT compound. After washing, permeabilization was performed using a permeabilization buffer (prepared by adding 10 μL of 0.5% Triton-X-100 to 1 mL of PBS) for 10 minutes. And sections were blocked with PBS containing 10% goat serum (#AR1009, Boster) for 1.5 hours at room temperature to reduce nonspecific binding. The samples were then incubated overnight at 4°C with primary antibodies against α-smooth muscle actin (α-SMA; #14395-1-AP, Proteintech) and CD68 (#MCA1957GA, Bio-Rad), each diluted 1:100. After incubation with primary antibodies, sections were washed three times with PBS and then incubated for 1 hour at room temperature with Alexa Fluor–conjugated secondary antibodies (goat anti-rabbit 546, #A11035, and goat anti-rat 488, #A11006; Invitrogen) at a 1:1,000 dilution. Nuclei were counterstained with DAPI (#C1005, Beyotime) for 30 minutes. Fluorescence images were acquired using a Leica TCS SP8 X confocal microscope, and signal intensity was measured with ImageJ (NIH, USA)^[13]^.

### Enzyme-Linked Immunosorbent Assay (ELISA)

To assess systemic inflammatory markers, serum concentrations of C-reactive protein (CRP; #KE10128, Proteintech) and serum amyloid A1 (SAA1; #CSB-EL020656MO, CUSABIO) were determined using specific ELISA kits following the manufacturers’ instructions. Briefly, serum samples and standards were added to antibody-coated wells, followed by incubation with the corresponding detection antibody and enzyme conjugate. After the colorimetric reaction was developed with the substrate solution, absorbance was recorded using a microplate reader (Rayto, Shenzhen, China). The final concentrations were calculated from standard calibration curves.

### TUNEL Staining

Apoptotic cells in the liver were detected on frozen liver sections using a One-Step TUNEL Apoptosis Detection Kit (#C1086, Beyotime) following the manufacturer’s protocol. The stained sections were imaged with a confocal microscope, and positive signals were quantified using ImageJ.

### RNA Sequencing

Total RNA was extracted from liver tissues, and mRNA was isolated using oligo(dT)-conjugated magnetic beads. The purified mRNA was fragmented at high temperature and used as a template for first-strand cDNA synthesis with reverse transcriptase. During the second-strand synthesis, end repair and A-tailing were performed, followed by adaptor ligation. The resulting cDNA fragments were purified with Hieff NGS® DNA Selection Beads to obtain target-sized fragments. Subsequently, PCR amplification was carried out to construct sequencing libraries. The quality of the libraries was assessed, and paired-end sequencing was performed on Illumina NovaSeq X Plus platform^[14]^. The original data files of RNA sequencing will be deposited on the Gene Expression Omnibus.

### Statistical Analysis

Data are presented as the mean ± standard deviation (SD) for normally distributed variables or as the median with interquartile range (IQR) for non-normally distributed variables. Normality of the datasets was assessed using the Shapiro–Wilk test. For two-group comparisons, normally distributed data were analyzed using a two-tailed unpaired Student’s *t*-test. When unequal variances were detected by the F-test, Welch’s correction was applied. Non-normally distributed data were analyzed with the two-tailed Mann–Whitney *U* test. For comparisons among more than two groups, data that were normally distributed but exhibited unequal variances were analyzed using the Brown-Forsythe ANOVA test, whereas non-normally distributed data were evaluated with the Kruskal-Wallis test. Statistical analyses were performed using GraphPad Prism 9.0 (GraphPad Software, San Diego, CA, USA), and *p* < 0.05 was considered statistically significant.

## Results and discussion

We first analyzed hepatic transcriptome data from patients at different stages of MASLD progression (GSE213621). Under physiological conditions, hepatic *THR-β* expression was approximately 4.5 times higher than that of *THR-α*. Notably, *THR-β* expression progressively declined with disease advancement, whereas *THR-α* expression remained relatively stable across different stages of MASLD (Fig. 1A). These observations indicate that alterations in *THR-β* expression accompany MASLD progression, providing a molecular context for further investigation of THR-β– targeting interventions such as Resmetirom in MASH.

**Figure 1.**
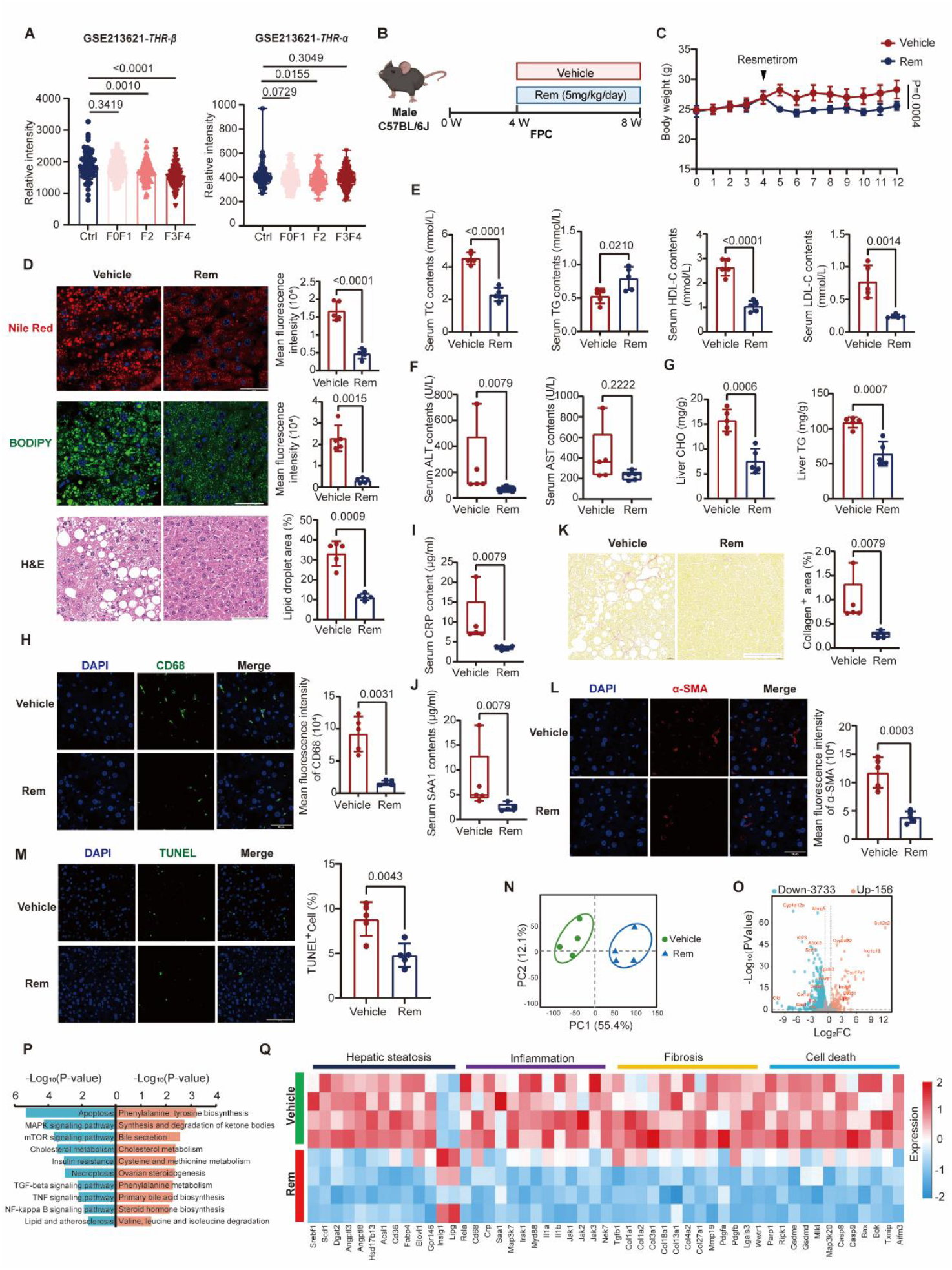
Therapeutic effects of Resmetirom on FPC diet–induced MASH. **(A)** Expression of thyroid hormone receptor-β (*THR-β*) and thyroid hormone receptor-α (*THR-α*) in human liver tissue across MASLD progression stages (GSE213621). **(B)** Experimental design: male C57BL/6J mice were fed a fructose, palmitate, and cholesterol (FPC) diet for 12 weeks. Therapeutic Resmetirom (Rem) (5 mg/kg/day) or vehicle (0.5% carboxymethyl cellulose-sodium + 0.4% Tween-80) was administered intraperitoneally from week 4 to week 12. **(C)** Body weight changes over the 12-week period. **(D)** Representative images and quantification of hepatic lipid accumulation by Nile Red, BODIPY, and hematoxylin & eosin (H&E) staining. Scale bar: 50 μm for Nile Red and BODIPY; Scale bar: 100 μm for H&E. **(E–F)** Serum biochemical analyses, including total cholesterol (TC), triglycerides (TG), high-density-lipoprotein-cholesterol (HDL-C), low-density lipoprotein-cholesterol (LDL-C), alanine aminotransferase (ALT) and aspartate aminotransferase (AST). **(G)** Total cholesterol and triglyceride content in liver. **(H)** Immunofluorescence analysis of macrophage infiltration (CD68). Scale bar: 50 μm. **(I-J)** Serum levels of systemic inflammation markers (CRP and SAA1). **(K–L)** Sirius Red and immunofluorescence staining of α-smooth muscle actin (α-SMA) indicating fibrosis. Scale bar: 100 μm for Sirius Red; Scale bar: 50 μm for α-SMA. **(M)** TUNEL assay for hepatocyte apoptosis. Scale bar: 100 μm. **(N)** Principal component analysis (PCA) of hepatic transcriptome showing distinct clustering between vehicle- and Resmetirom-treated groups (n=4). **(O)** Volcano plot depicting differentially expressed genes (DEGs) between groups (p < 0.05, |log_2_FC| > 0.5). **(P)** Pathway enrichment analysis of DEGs highlighting key biological processes. **(Q)** Heatmap of representative DEGs associated with hepatic steatosis, inflammation, fibrosis, and cell death. Statistical significance: *p* < 0.05.

To further investigate the mechanism of Resmetirom’s action, we established a human-relevant diet-induced MASH model using a fructose, palmitate, and cholesterol (FPC) diet^[11]^. Mice were maintained on the FPC diet for 12 weeks, with therapeutic administration of Resmetirom (5 mg/kg/day) initiated after 4 weeks of feeding and continued for an additional 8 weeks via intraperitoneal injection. The vehicle consisted of 0.5% carboxymethyl cellulose-sodium (CMC-Na) and 0.4% Tween-80 (Fig. 1B).

Resmetirom treatment resulted in lower body weight, accompanied by marked improvement of hepatic steatosis as shown by Nile Red, BODIPY, and H&E staining (Fig. 1C, D). Serum biochemical analyses revealed significantly lower levels of total cholesterol (TC), high-density-lipoprotein-cholesterol (HDL-C), low-density-lipoprotein-cholesterol (LDL-C), and alanine aminotransferase (ALT), accompanied by a slight increase in triglycerides (TG) (Fig. 1E, F). Hepatic TG and TC contents were also diminished following Resmetirom treatment (Fig. 1G). Immunohistochemical staining revealed a marked decrease in CD68 expression in the liver, indicating reduced macrophage infiltration (Fig. 1H). Notably, we observed for the first time that Resmetirom treatment significantly lower serum levels of C-reactive protein (CRP) and serum amyloid A1 (SAA1), two liver-derived biomarkers widely used to assess systemic inflammation, underscoring its potential to attenuate inflammatory responses (Fig. 1I, J). This anti-inflammatory effects of Resmetirom also have clinical implications as chronic inflammation is a common driving mechanism for pan-vascular and pan-metabolic diseases ^[15-18]^. Sirius Red staining revealed decreased collagen deposition, while immunofluorescence staining also demonstrated reduced α-SMA expression, further indicating attenuation of hepatic fibrosis (Fig. 1K, L). Consistently, TUNEL staining demonstrated a marked decline in cell death in the liver of Resmetirom-treated mice. (Fig. 1M). Collectively, these results demonstrate that Resmetirom effectively ameliorates hepatic steatosis, inflammation, fibrosis, and liver injury in FPC diet–induced MASH.

To identify genome-wide transcriptional programs modulated by Resmetirom and thus pinpoint candidate pathways mediating its hepatoprotective effects, we performed bulk RNA sequencing of liver tissues from vehicle- and Resmetirom-treated mice. Principal component analysis (PCA) of variance-stabilized expression values showed clear separation of treatment groups, indicating that Resmetirom induced a robust transcriptional response (Fig. 1N). Using the criteria of *p* < 0.05 and |log_2_ (fold change, FC) | > 0.5, a total of 3,733 genes were downregulated and 173 genes were upregulated (Fig. 1O). Functional enrichment analysis showed that these differentially expressed genes (DEGs) were predominantly associated with pathways involved in inflammation, cholesterol metabolism, and apoptosis (Fig. 1P). To gain insight into the molecular alterations underlying Resmetirom’s effects, we visualized representative DEGs in a heatmap. The expression patterns indicated enrichment in pathways associated with steatosis, inflammation, fibrosis, and cell death (Fig. 1Q). Genes involved in lipid metabolism, such as sterol regulatory element-binding transcription factor 1 (*Srebf1*) and stearoyl-CoA desaturase 1 (*Scd1*), as well as inflammatory genes including interleukin-1 alpha (*IL-1α*), interleukin-1 beta (*IL-1β*), and fibrosis-related genes such as collagen type I alpha 1 (*Col1α1*) and collagen type I alpha 2 (*Col1α2*), were significantly downregulated in the Resmetirom-treated group (Fig. 1Q).

Importantly, multiple genes implicated in distinct forms of programmed cell death were markedly downregulated in the Resmetirom-treated group (Fig. 1Q). Specifically, poly (ADP-ribose) polymerase 1 (*Parp1*), receptor-interacting serine/threonine-protein kinase 1 (*Ripk1*), mixed lineage kinase domain-like pseudokinase (*Mlkl*), and mitogen-activated protein kinase kinase kinase 20 (*Map3k20*) are central mediators of apoptosis, necroptosis and ribotoxic stress response signaling. Gasdermin D (*Gsdmd*) and gasdermin E (*Gsdme*) act as pore-forming executors of pyroptosis, while caspase-8 (*Casp8*) and caspase-9 (*Casp9*) serve as key initiator caspases driving extrinsic and intrinsic apoptosis, respectively. In addition, BCL-2 antagonist/killer 1 (*Bak*), BCL-2-related ovarian killer (*Bok*), thioredoxin-interacting protein (*Txnip*), and apoptosis-inducing factor mitochondria-associated 3 (*Aifm3*) participate in mitochondrial- and oxidative stress–associated caspase-independent death pathways. The coordinated downregulation of these genes suggests that Resmetirom may confer hepatoprotection not only through improving lipid metabolism and inflammatory responses, but also by broadly repressing hepatocyte cell death programs that contribute to MASH progression. These findings underscore a previously underappreciated role of THR-β activation in modulating hepatic cell death; a process increasingly appreciated as a key contributor to MASH progression^[19, 20]^.

In summary, the present study demonstrates that Resmetirom exerts multifaceted protective effects against MASH progression by suppressing hepatic lipid accumulation, inflammation, fibrosis, and cell death. These findings provide new mechanistic insights into THR-β-mediated hepatoprotection and suggest that targeting cell death may represent an important therapeutic dimension of Resmetirom’s clinical efficacy in MASH which warrants clinical investigation.

## Acknowledgements

This study was supported by the National Natural Science Foundation of China (Nos. 82370444 and 12411530127). Additional support was provided by the Program for Innovative Research Team of The First Affiliated Hospital of USTC (CXGG02), the USTC Research Funds of the Double First-Class Initiative (No. YD9110002089), and the Research Funds of the Centre for Leading Medicine and Advanced Technologies of IHM (2025IHM01040).

## Data Availability

The public datasets used in this study include GSE213621 (different stages of MASLD).

## Author contributions

Suowen Xu and Jianping Weng conceptualized the project. Chenyang He performed the experiments, analyzed the data, and wrote the manuscript. Suowen Xu and Jianping Weng provided conceptual inputs and suggestions into the manuscript. Zhihua Wang assists in bioinformatic analysis of RNA-seq data.

